# Combining feature selection and shape analysis uncovers precise rules for miRNA regulation in Huntington’s disease mice

**DOI:** 10.1101/2020.01.24.918540

**Authors:** Lucile Mégret, Satish Sasidharan Nair, Julia Dancourt, Jeff Aaronson, Jim Rosinski, Christian Neri

## Abstract

**Background:** MicroRNA (miRNA) regulation is associated with several diseases, including neurodegenerative diseases. Several approaches can be used for modeling miRNA regulation. However, their precision may be limited for analyzing multidimensional data. Here, we addressed this question by integrating shape analysis and feature selection into miRAMINT, a methodology that we used for analyzing multidimensional RNA-seq and proteomic data from a knock-in mouse model (Hdh mice) of Huntington’s disease (HD), a disease caused by CAG repeat expansion in huntingtin (htt). This dataset covers 6 CAG repeat alleles and 3 age points in the striatum and cortex of Hdh mice.

**Results:** Remarkably, compared to previous analyzes of this multidimensional dataset, the miRAMINT approach retained only 31 explanatory striatal miRNA-mRNA pairs that are precisely associated with the shape of CAG repeat dependence over time, among which 5 pairs with a strong change of target expression levels. Several of these pairs were previously associated with neuronal homeostasis or HD pathogenesis, or both. Such miRNA-mRNA pairs were not detected in cortex.

**Conclusions:** These data suggest that miRNA regulation has a limited global role in HD while providing accurately-selected miRNA-target pairs to study how the brain may compute molecular responses to HD over time. These data also provide a methodological framework for researchers to explore how shape analysis can enhance multidimensional data analytics in biology and disease.

## Background

Several neurodegenerative diseases (NDs) such as Alzheimer’s disease, Parkinson’s disease, Amyotrophic lateral sclerosis and Huntington’s disease (HD) may evolve through gene deregulation, which has fostered a large number of studies aiming to explore the role of micro-RNA (miRNA) regulation in driving gene deregulation in these diseases [1–5]. MiRNAs are short (~21nt) non-coding RNAs that regulate gene expression through the degradation or translational repression of mRNAs. Although miRNAs are believed to play a discrete as well as global role in NDs such as HD [3, 6–8], the identification of miRNAs that on a system level could be central to ND pathogenesis remains challenging [3]. Part of this problem relates to the lack of rich data, *e.g.* time series data, or sufficiently homogeneous data, *e.g.* in tissues and subjects [1]. This problem also relates to the challenges associated with accurately modeling miRNA data and mRNA data on a system level. To this end, several approaches predict miRNA targets based on binding sites, where the most commonly used features for predicting miRNA targets include sequence complementarity between the “seed” region of a miRNA and the “seed match” region of a putative target mRNA, species conservation, thermodynamic stability and site accessibility [9]. These methods can be classified in two categories. One category comprises heuristic methods [10] such as for example TargetScan [11] and mirSVR [12]. However, the number of possible targets for a single miRNA can be large, greatly limiting biological precision. The other category comprises machine-learning techniques (*e.g.* decision trees, support vector machine and artificial neural networks) such as mirMark [9], TarPmiR [13], TargetMiner [14], TargetSpy [15] and MiRANN [16]. More sophisticated algorithms in this category of methods include deep learning methods such as for example DeepMirTar [17]. Finally, this category also comprises combinatorial ensemble approaches for improving the coverage and robustness of miRNA target prediction [18].

Besides predicting binding sites, another strategy for predicting miRNA targets is to search for negative correlations between miRNA and target expression levels. Such approaches include the use of Bayesian analysis such as GeneMiR++ [19]. However, optimal fitting between miRNAs and putative targets upon Bayesian causal inference can be biased due to building a large and heterogenous network of causal interactions that involves miRNA-to-miRNA, target-to-target and target-to-miRNA interactions in addition to miRNA-target interactions [20]. To overcome this problem, Bayesian models may be filtered using external database information on miRNA binding sites [21]. However, filtering does not address the problem of miRNA effect sizes nor takes into account the possibility that miRNA-target interactions could be indirect eventhough there is evidence for a binding site in external databases. Expression-based approaches also involve support vector machine analysis [22], Gaussian process regression model [23] and network inference such as weighted gene correlation network analysis (WGCNA), the latter approach which has been used, for example, for modelling miRNA regulation in hepatitis C [24] and in HD knock-in mice (Hdh mice) [3].

Although network inference methods such as Bayesian analysis and WGCNA may provide insights into the features of miRNA regulation, they may be prone to aggregation of a large number of hypotheses around strongly deregulated entities [3, 20], lacking discriminative power and biological precision, and impairing data prioritization. Here, we addressed this problem by developing an approach in which network-based analysis for reducing data complexity is followed by robust random-forest (RF) analysis for selecting explanatory variables (*i.e.* miRNAs best explaining targets, with a *P*-value computed for each predictor variable and each predictor variable stable across RF iterations involving different seeds) and shape analysis (surface matching) for building discriminative and accurate ensembles of negatively correlated miRNA-mRNA pairs. We used RF analysis for feature selection as this method does not make any prior hypothesis on the existence of a relationship, whether direct or indirect, between a miRNA and a target. To select the most interesting miRNAs, this analysis was supplemented with evidence for binding sites as instructed from multiple databases and followed by data prioritization using criteria such as CAG-repeat-length dependence and the fold change of target expression. We applied this approach to the analysis of multidimensional data in the allelic series HD knock-in mice (Hdh mice), currently the largest and more comprehensive datasets (6 CAG-repeat lengths, three age points, several brain areas: miRNA, mRNA and proteomic data) to understand how miRNA regulation may work on a system level in neurodegenerative diseases [2]. We focused on the study of miRNA regulation mediated by mRNA degradation as the coverage and dynamics of proteomic data in the allelic series of Hdh mice is limited compared to miRNA and mRNA data. As developed below, we found that, on a global level, miRNA data explains a very small proportion of the CAG-repeat- and age-dependent dynamics of gene deregulation in the striatum (and none in cortex) of Hdh mice, retaining 31 miRNA-mRNA pairs implicated in neuronal activity and cellular homeostasis, among which only five pairs are of high interest.

## Results

### Multimodal selection of miRNA targets

To understand how the dynamics of miRNA regulation may work on a system level in the brain of Hdh mice, we applied miRNA regulation analysis via multimodal integration (miRAMINT), a pipeline in which novelty is to combine shape analysis with random forest analysis (Figure 1).

**Figure 1.**
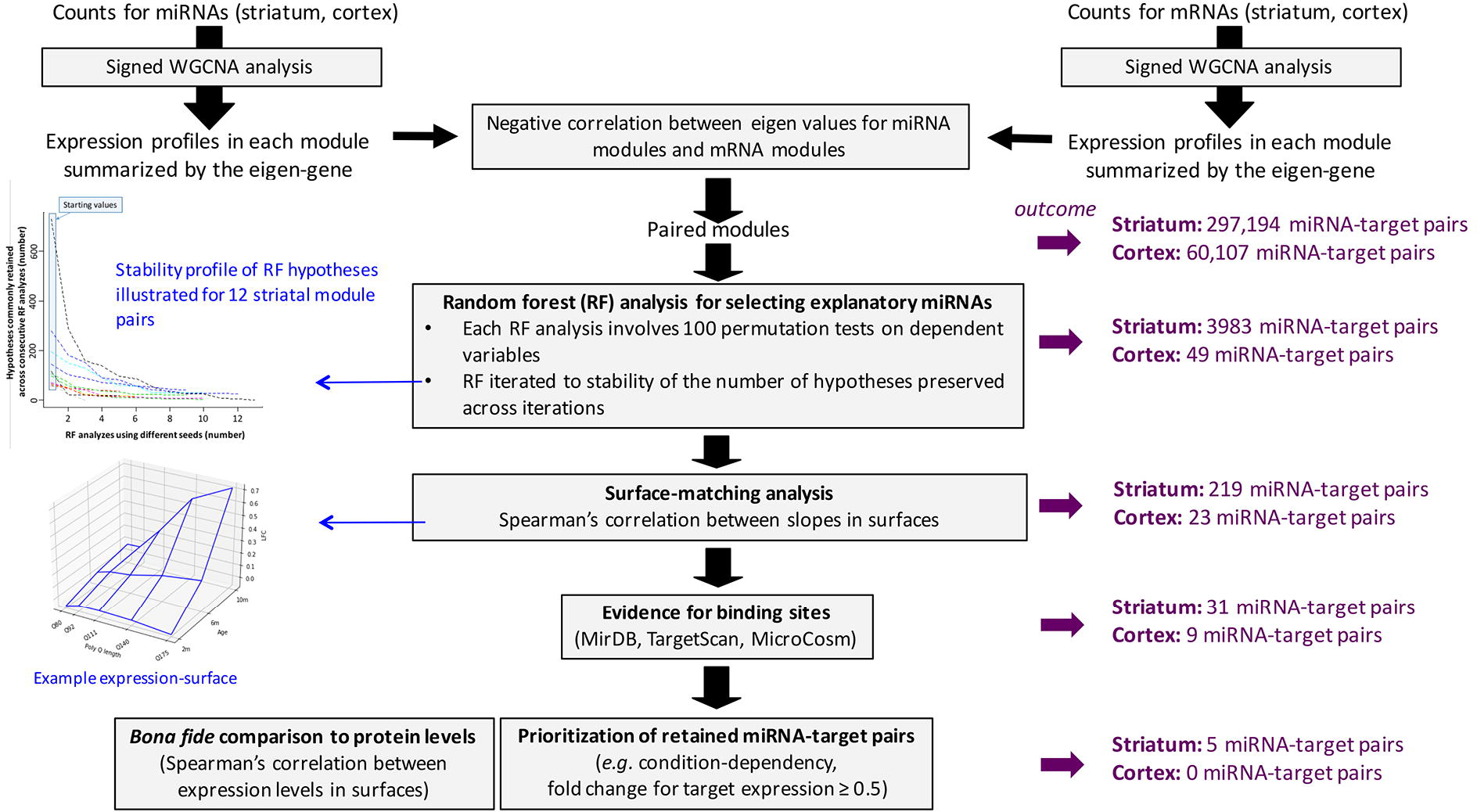
MiRAMINT analysis of miRNA regulation. This protocol integrates network-based analysis for reduction of data complexity followed by (i) random forest (RF) analysis for selecting explanatory variables, with a p-value computed for each predictor variable using the Altmann’s approach [29] and this RF analysis iterated (which involves different seeds in each iteration) until the number of hypotheses is stable across consecutive iterations (see also Materials and Methods), (ii) surface-matching analysis for high precision in matching the miRNA and mRNA expression profiles across conditions (herein as defined by 6 CAG repeat alleles and 3 age points), (iii) evidence for binding sites and (iv) data prioritization for selecting predicted miRNA-mRNA pairs of high interest. The number of possible miRNA-mRNA pairs retained at each step of the analysis (outcome) of multidimensional data from the brain of Hdh mice is indicated. The miRNA-mRNA pairs retained upon shape-matching analysis can be visualized at http://www.broca.inserm.fr/MiRAMINT/index.php. The whole approach, data prioritization included, retained 5 miRNA-mRNA pairs of high interest in the striatum of Hdh mice and none in the cortex.

As a first step, we performed a signed WGCNA analysis [25] of mRNA and miRNA expression profiles to reduce data complexity through building co-expression modules. The expression profiles of genes (respectively miRNA) in each cluster were summarized using the eigen-gene (respectively eigen-miRNA) [26]. We then selected the miRNA module(s) where the eigen-miRNAs are negatively correlated with the eigen-genes. This analysis retained 8 miRNA co-expression modules and 18 target co-expression modules in the striatum and 4 miRNA co-expression modules and 14 gene co-expression modules in the cortex (Table S1, see http://www.broca.inserm.fr/MiRAMINT/index.php for edge lists). Amongst all possible associations (144) between miRNA modules and target modules, 12 negative correlations (false discovery rate lower than 1%) were retained in the striatum and in the cortex (Table 1).

**Table 1.** Negative correlations (FDR < 0.01) between miRNA and mRNA modules in Hdh mice.

| Striatal data |  | Cortical data |  |
| --- | --- | --- | --- |
| miRNA module | Target module | miRNA module | Target module identity |
| *ID#1 (N= 206) | ID#2 (N=40131) | ID#1 (N=217) | ID#1 (N=7028) |
| ID#1 (N= 206) | ID#7 (N=108) | ID#1 (N=217) | ID#3 (N=325) |
| ID#1 (N= 206) | ID#11 (N=49) | ID#1 (N=217) | ID#6 (N=223) |
| ID#1 (N= 206) | ID#15 (N=41) | ID#2 (N=149) | ID#11 (N=66) |
| ID#1 (N= 206) | ID#17 (N=33) | ID#3 (N=87) | ID#4 (N=291) |
| ID#2 (N= 138) | ID#18 (N=27) | ID#3 (N=87) | ID#8 (N=170) |
| ID#3 (N= 104) | ID#5 (N=163) | ID#3 (N=87) | ID#9 (N=75) |
| ID#3 (N= 104) | ID#6 (N=160) | ID#3 (N=87) | ID#10 (N=74) |
| ID#3 (N= 104) | ID#10 (N=51) | ID#4 (N=82) | ID#4 (N=291) |
| ID#3 (N= 104) | ID#11 (N=49) | ID#4 (N=82) | ID#8 (N=170) |
| ID#4 (N= 63) | ID#3 (N=935) | ID#4 (N=82) | ID#9 (N=75) |
| ID#8 (N= 20) | ID#2 (N=40131) | ID#4 (N=82) | ID#10 (N=74) |

\*Identity number.

We then tested whether the log fold change (LFC) for miRNA expression across the 15 CAG-repeat and age-dependent conditions tested in Hdh mice might explain target expression levels across these conditions. To this end, we applied RF analysis, which allows this question to be addressed in an unbiased manner (*i.e.* with no *a priori* hypothesis on the existence of miRNA-target relationships) and which has been successfully used to study miRNA regulation on a binding site level [27, 28]. To ensure a strong level of reliability, we applied a version of RF analysis in which a *P*-value (based on 100 permutations) is computed for each predictor variable using the Altmann’s approach [29] and in which each hypothesis on a predictor variable is stable across RF iterations involving different seeds (See Materials and Methods). This approach retained 3983 pairs (involving 141 explanatory miRNA variables and 350 dependent genes variables) in the striatum and 49 pairs (involving 16 explanatory miRNA variables and 3 dependent genes variables) in the cortex (Table S2). Next, we tested whether the shape of the surface defined by the LFC values for explanatory miRNAs is negatively correlated with that defined by the LFC values for the corresponding targets (see Methods). Surface-matching retained 219/3983 relationships in the striatum, and 23/49 relationships in the cortex (Table S2). Finally, in these latter groups of miRNA-target relationships, we retained those showing evidence for binding sites as indicated in the TargetScan [11], MicroCosm [30] and miRDB [31] databases, which generated a final number of 31 predictions (14 miRNAs explaining 20 targets) in the striatum and 9 predictions (6 miRNAs explaining 3 targets) in the cortex (Table S2). No overlap was found with miRTarBase, a database which contains experimentally-validated miRNA-mRNA pairs. Thus, remarkably, integrating shapes and random forests in miRAMINT selected quite a small number of miRNA-target pairs that show significant htt- and age-dependent features in the brain of Hdh mice.

### Comparison to *bona fide* information contained in proteomic data

Gene and protein expression data from the same cells under similar conditions usually do not show a strong positive correlation [32–35]. As shown above, miRAMINT is a selective data analysis work-flow in which a small number of htt- and time-dependent miRNA regulation events may be retained, thus reducing the expectation for changes in protein expression levels to be correlated with changes in corresponding open reading frames. Nonetheless, we assessed whether some of the dynamics of gene deregulation explained by the dynamics of miRNA expression in the brain of Hdh mice might be associated with comparable dynamic changes of protein levels. To this end, we focused on the striatal miRNA-target pairs identified in the striatum as the brain area where gene deregulation is the strongest [2] and where miRNA levels are reliably associated with mRNA levels by miRAMINT, which represents 20 targets (Table S2). We observed that 9/20 targets (45%) retained by miRAMINT have at least one corresponding protein, from which only 3 targets (15%) were positively correlated with protein products across CAG repeat lengths and age points (Table S3). Although this overlap is limited, these observations provided *bona fide* information for data prioritization as developed below.

### Data prioritization upon miRAMINT analysis

Although selective, data analysis in miRAMINT enables a diversity of profiles in terms of CAG-repeat dependence, age dependence and magnitude of effects across conditions to be retained. Several criteria may then be used for prioritizing the most interesting pairs, including (i) the overall shape of the gene deregulation plane (*e.g.* linear effects, biphasic effects, local effects) and the maximal amplitude of gene deregulation at any point in the CAG repeat- and age-dependent plane, (ii) the strength of plane matching (*i.e.* the Spearman’s score for surface-matching), (iii) the number of databases concluding to a binding site between miRNA(s) and predicted target(s) and (iv), if available, positive correlations between changes in the expression of proteins and of genes encoding these proteins.

The analysis retained 31 miRNA-mRNA pairs in the striatum, among which 17 top pairs corresponding to either binding sites found in more than one miRNA target database or highest Spearman’s score for surface matching, or both (Figure 2A), including 5 pairs for which from the maximally-achieved log fold change of target is greater than or equal to 0.5 (Figure 2B). Biological annotations suggested this group of miRNA-target pairs may be notably implicated in Jak-STAT signaling, Th1 and Th2 cell differentiation, ether lipid metabolism and N-glycan biosynthesis signaling pathway (Figure 2A).

**Figure 2.**
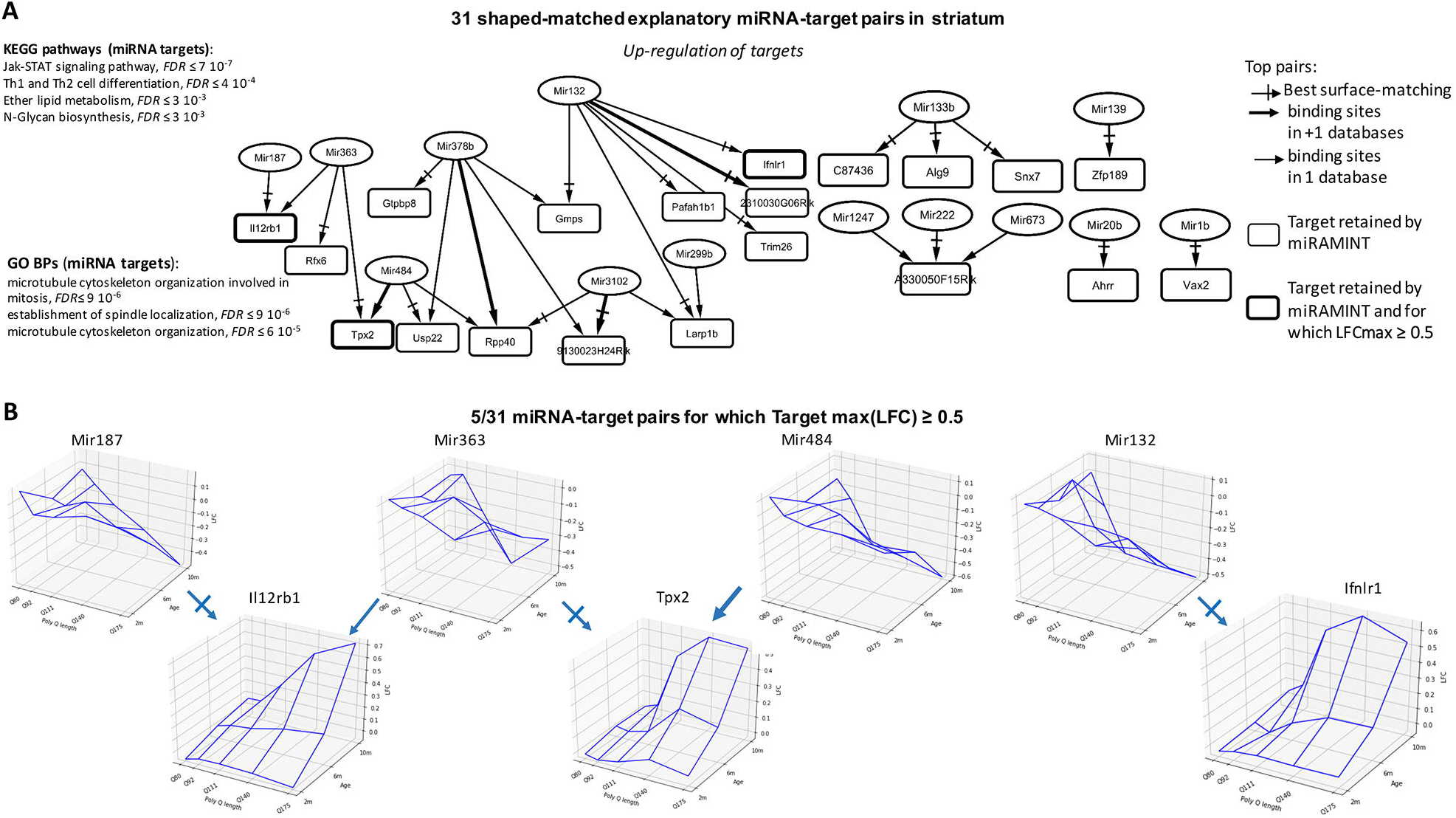
Selected miRNA-target pairs in the striatum of Hdh mice. (**A**) Shown are the 31 miRAMINT miRNA-target pairs (see also Table S3 for the full list of miRAMINT miRNA-target pairs in striatum). The targets are contained in rectangles and the miRNAs in ellipses. A thick edge means that evidence for binding sites is available from at least two miRNA databases. A thin edge means that evidence for binding sites is available from only one miRNA database. A thick rectangle means that the maximal LFC of the target is greater than 0.5. A cross arrow indicates the miRNA that is best paired with a target when this target has several possible miRNA regulators. Biological annotations of miRNA targets correspond to GO Biological processes or KEGG pathways at the result of STRING analyzes using stringent criteria (*i.e.* STRING score > 0.7, Databases and Experiments only, 20 neighbors added on the first shell) the KEGG pathways are those with, at least, 3 genes implied, the GO Biological processes are those with, at least, 5 genes implied. (**B**) Examples of 3D-graphs for top miRNA-target pairs (LFC amplitude of the target above 0.5).

In the cortex, miRAMINT retained 9 miRNA-target pairs that tend to show a biphasic (deregulation at 6 months, then return to initial level) age-dependent profile, including 6 miRNAs and 3 targets annotated for inflammatory pathways (Tnfrs11a) such as NF-kappa B signaling, a pathway involved in neuronal apoptosis [36], and for cell genesis and death (protogenin, cadherin 9) (Figure 3). However, deregulation in these miRNA-target pairs was not dependent on the CAG repeat lengths in a strongly consistent (linear effect) manner, contrasting with the consistency for CAG repeat dependence in the striatum (Figure 2B). Additionally, raising the threshold on the log fold change of target expression to a value of 0.5 reduced the number of top predictions to 0 in the cortex. Thus, miRAMINT analysis indicated that no miRNA-target pairs are consistently and strongly deregulated in a CAG-repeat- and age-dependent manner in the cortex of Hdh mice.

**Figure 3.**
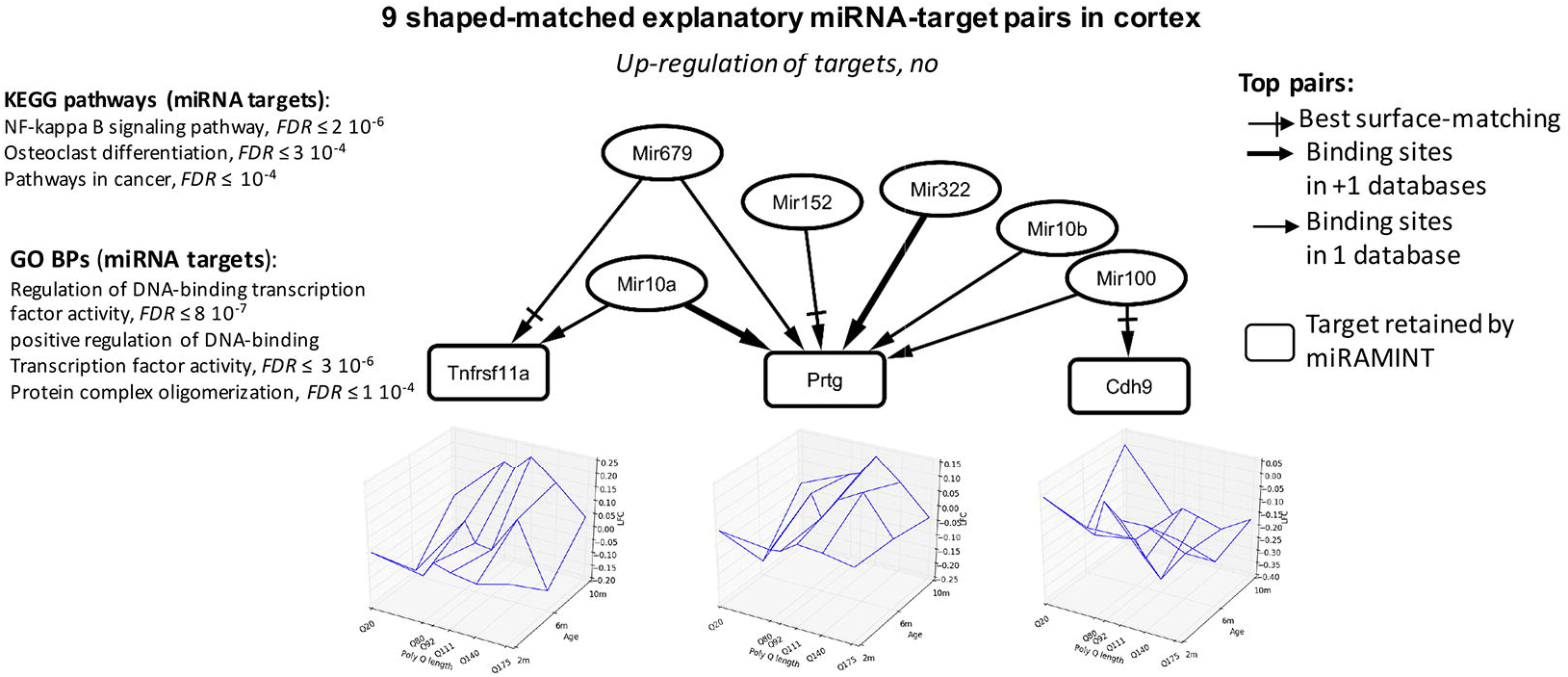
Selected miRNA-target pairs in the cortex of Hdh mice. Shown are the 9 miRAMINT miRNA-target pairs (see also Table S3 for the full list of miRAMINT miRNA-target pairs in cortex). The targets are contained in rectangles and the miRNAs in ellipses. A thick edge means that evidence for binding sites is available from at least two miRNA databases. A thin edge means that evidence for binding sites is available from only one miRNA database. All LFC are below 0.5. A cross arrow indicates the miRNA that is best paired with a target when this target has several possible miRNA regulators. Biological annotations of miRNA targets correspond to GO Biological processes or KEGG pathways at the result of STRING analyzes using stringent criteria (*i.e.* STRING score > 0.7, Databases and Experiments only, 5 neighbors added on the first shell) the KEGG pathways are those with, at

## Discussion

As multi-point data become available for modeling miRNA regulation [2], comprehensive approaches are needed to build precise models of miRNA regulation of gene expression. Here, we addressed this problem by integrating several machine learning concepts, each of them bringing complementary elements of information and reliability about the way that miRNA levels and target levels may evolve across conditions. MiRAMINT analysis (Figure 1) comprises WGCNA analysis for reducing data complexity, followed by (i) RF analysis for selecting explanatory variables, in which a p-value is computed for each predictor variable and in which RF analysis is iterated (involving different seeds) until the number of hypotheses is stable across consecutive iterations, (ii) shape analysis for matching the miRNA and mRNA expression profiles across conditions, (iii) evidence for binding sites and (iv) *bona fide* comparison of the gene targets retained into the model to protein expression profiles.

Since the coverage and dynamics of proteomic data in the allelic series of Hdh mice are limited compared to those of miRNA and mRNA data, we focused our study on modeling miRNA regulation mediated by mRNA degradation. Depending on the features of input data layers, miRAMINT analysis may be used to analyze gene expression repression mediated by mRNA degradation or inhibition of protein translation, or both.

Combining shape analysis and feature selection for negatively correlating miRNA and mRNA data suggests that miRNA regulation via mRNA degradation may have a limited global role in the striatum and cortex of Hdh mice. This conclusion is supported by the small number of miRNA-target relationships that show a consistent pattern (*i.e.* strong and linear effects) of expression in the surface defined by CAG-repeat lengths and age points in the striatum of these mice. This conclusion is reminiscent of a similar trend detected in the brain of wild-type mice, where miRNA regulation may be poorly correlated to gene expression signatures across cell types [38]. This conclusion is even more stringent for the cortex of Hdh mice, suggesting that miRNA regulation do not play a critical role in truly responding to HD in this brain area. In so far, our model significantly differs from a previous analysis [3] of the RNA-seq time series data in the allelic series of Hdh mice [2] in which global (eigenvalue-based) negative correlation between miRNAs and target modules (using WGCNA) was used to build a model of miRNA regulation. Although some of the miRNAs retained by miRAMINT analysis were also retained in this former study [3] (see Table S3: 12/14 miRNAs common to the two studies), miRAMINT miRNA-target pairs are in smaller numbers (before data prioritization: 31 miRAMINT predictions in striatum, instead of 7514 WGCNA predictions contained in 55 negative correlations between miRNA and target modules in striatum; 9 miRAMINT predictions in cortex, instead of 186 WGCNA-based predictions contained in 9 negative correlations between miRNA and target modules) and, importantly, except to one case (Mir132-Pafah1b1), they are associated with different targets. These differences are likely due to the higher accuracy associated with tree-based analysis combined with surface matching in miRAMINT compared to using a global (eigenvalue-based) negative correlation scheme between target modules and miRNAs [3].

A former bioinformatic analysis of miRNA expression identified 33 possible miRNA-target relationships in *post-mortem* brain samples of HD patients compared to control individuals [37]. We found no overlap between these predictions and the miRNA-target pairs retained by miRAMINT, which is expected as the study of *post-mortem* brain samples relied on a simple overlap analysis (based on binding sites in TargetScan) between lists of differentially expressed miRNAs and mRNAs [39] and as miRNA regulation in the humain brain could significantly differ from that in the mouse brain.

The lack of miRNA-target pairs that may truly function in a CAG-repeat dependent manner in the cortex of Hdh mice is intriguing. Although some of the miRNAs retained in our analysis showed age- and CAG-repeat-dependent profiles, all nine miRNA-target pairs (involving 3 targets) show a bi-phasic response with deregulation at 6 months of age and return to initial (2-month) expression levels at 10 months of age. Since miRNA regulation may be highly dependent on cellular context, we speculate this observation could relate to the large heterogeneity of neuronal populations in cortex, which could preclude a sufficiently sensitive analysis of HD and age-dependent miRNA regulation in whole cortex extracts compared to whole striatum extracts. Alternatively, this observation could relate to a strong level of miRNA-regulation reprogramming and impairment in the HD cortex, as further discussed below.

Although we cannot exclude the possibility that the conclusion about a limited global role of miRNA regulation in the brain of Hdh mice might be biased by the current lack of cell-type specific RNA-seq data in HD mice, our data highlight a new set of precisely matched and highly prioritized miRNA-target relationships (see Figure 2, Table S3) that are known to play a role in neuronal activity and homeostasis. This feature applies to miRNAs that are upregulated in the striatum of Hdh mice. Mir132 (upregulated and paired with 2310030G06Rik, the Guanine Monophosphate Synthase Gmps, Interferon Lambda Receptor Ifnlr1, Ribonucleoprotein Domain Family Member Larp1b, Platelet Activating Factor Acetylhydrolase 1b Regulatory Subunit Pafah1b1 and Tripartite Motif-Containing ProteinTrim26) is associated to brain vascular integrity in zebrafish [38], spine density [39] and synaptogenesis [40]. Knocking down Mir1b (upregulated and paired with Ventral Anterior Homeobox 2, Vax2) significantly alleviated neuronal death induced by hypoxia [41]. miR139 (paired with the zinc finger protein 189 Zfp189) modulates cortical neuronal migration by targeting Lis1 in a rat model of focal cortical dysplasia [42]. Mir20b (paired with the Aryl-Hydrocarbon Receptor Repressor Ahrr) inhibits cerebral ischemia-induced inflammation in rats [43]. Exosomes harvested from Mir133b (paired with C87436, alpha-1,2-mannosyltransferase Alg9 and sorting nexin Snx7) overexpressing mesenchymal stem cells may improve neural plasticity and functional recovery after stroke in the rat brain [44]. In addition, Mir133b may promote neurite outgrowth via targeting RhoA [45] and miR-133b may be critical for neural functional recovery after spinal cord injury and stroke in several organisms [46–48]. Mir187 (paired with the Interleukin 12 Receptor Subunit Beta Il12rb1) is associated with the regulation of the Potassium Channel KCNK10/TREK-2 in a rat epilepsy model [49]. Finally, Mir363 is involved in neurite outgrowth enhanced by electrical stimulation in rats [50]. Target genes retained by MiRAMINT analysis in the striatum are also relevant to neuronal activity and homeostasis. Usp22 (targeted by Mir484 and Mir378b) was previsouly implicated in the maintenance of neural stem/progenitor cells via the regulation of Hes1 in the developing mouse brain [51]. Trim26 is related to DNA damage repair and cellular resistance to oxidative stress [52, 53]. In addition, neuroinformatic analyses have linked Trim26 to neuropsychiatric disorders such as anxiety disorders, autistic spectrum disorders, bipolar disorder, major depressive disorder, and schizophrenia [54]. Tpx2 (targeted by Mir484 and Mir363), promotes acentrosomal microtubule nucleation in neurons [55] and regulates neuronal morphology through interaction with kinesin-5 [56]. During eye and brain neurogenesis, the Xvax2 protein was detected in proliferating neural progenitors and postmitotic differentiating cells in ventral regions of both structures in Xenopus embryos [57]. Snx7 has been related to Alzheimer’s disease pathogenesis through the reduction of amyloid-beta expression [58]. In addition, Snx7 may participate in the control of glutamatergic and dopaminergic neurotransmission via the regulation of the kynurenine pathway, which is related to psychotic symptoms and cognitive impairment [59]. Finally, Pafah1b1 (targeted by Mir132), has been associated with the abnormal migration of cortical neurons and with neurologic disorder in mice and humans [60, 61]. In cortex, very few miRNA-target pairs were retained, and they involve target genes with low-amplitude fold change of expression. Nonetheless, it is interesting to note that some of the miRNA retained in the cortex were associated with neuronal homeostasis. Mir10a (paired with the TNF receptor superfamily member Tnfrsf11a/RANK, involved in inflammatory response in the mouse [62], and with protogenin Prtg, involved in neurogenesis and apoptosis [63, 64]) and Mir10b (paired with protogenin Prtg) are associated with the modulation of brain cell migration and aging [65, 66]. MiRNA322 (paired with protogenin Prtg) is associated with apoptosis and Alzheimer’s disease (AD) [67]. Finally, Mir100 (paired with cadherin Cdh9), is associated with neurological disorders such as AD, schizophrenia and autism [68–71].

Since miRAMINT finely accounts for the disease- and time-dependent features of miRNA and mRNA data in Hdh mice, miRAMINT miRNA-target pairs are strongly relevant to how cells and tissues may compute responses to HD on a miRNA regulation level. Amongst the 14 miRNAs retained by MiRAMINT analysis in the striatum (see Figure 2A), it is interesting to note that the levels of Mir222 (paired with A330050F15Rik) are increased in the plasma of HD patients and, however, were reported to be decreased in the striatum of transgenic 12-month-old YAC128 and 10-week-old R6/2 mice [72, 73]. Here, our analysis puts forth the downregulation of Mir222 as an event that is highly CAG-repeat and age-dependent in Hdh mice and, therefore, that may be strongly relevant to the response of the mouse striatum to HD.

## Conclusions

In summary, we addressed the problem of accurately modeling the dynamics of miRNA regulation from the analysis of multidimensional data. Our study puts forth the added value of combining shape analysis with feature selection for predictive accuracy and biological precision in modeling miRNA regulation from complex datasets, as illustrated by precise self-organised learning from multidimensional data obtained in the striatum and cortex of HD knock-in mice. MiRAMINT provides a convenient framework for researchers to explore how combining shape analysis with feature selection can enhance the analysis of multidimensional data in precisely modeling the interplay between layers of molecular regulation in biology and disease.

## Methods

### Source data

The RNA-seq (mRNA and miRNA) data analyzed herein originate from the striatum and cortex of the allelic series of Hdh knock-in mice (Q20, Q80, Q92, Q111, Q140 and Q175 at 2, 6 and 10 months of age) as previously reported [2]. The GEO IDs for transcriptome profiling data in Hdh mice are GSE65769 (Cortex, miRNAs), GSE65773 (Striatum, miRNAs), GSE65770 (Cortex, mRNAs) and GSE65774 (Striatum, mRNAs).

### Conversion between gene symbols and Entrez identifiers

We consistently used Entrez identifiers to unambiguously identify genes. Since many external resources identify genes by symbols, we used the Bioconductor package (https://www.bioconductor.org/) to convert gene symbols to Entrez identifiers. Typically, not all gene symbols can be unambiguously mapped to an Entrez ID; genes with such ambiguous mappings were kept with the Entrez identifiers.

### Removal of outliers in expression data

To remove outliers, we used variance stabilization to transform counts. Within each tissue and for each age-point, we constructed an Euclidean-distance sample network and removed those samples whose standardized inter-sample connectivity Z.k was below a threshold set to 2.5.

### Differential expression analysis

mRNA and miRNA significant read-count data for eight individuals (four males and four females) as available in the RNA-seq data in the allelic series of Hdh mice was fed into Deseq2 implemented in the R package DESeq2 [24] in order to obtain a log-fold-change (LFC) vector for each condition (CAG-repeat length, age) and a vector indicating if the genes are up-regulated (LFC > 0 and p-value < 0.05), down-regulated (LFC < 0 and p-value < 0.05) or unchanged (p-value ≥ 0) for each condition. The set age is k, and Q20 was used as reference for each condition at age k and Q>20.

### MiRAMINT analysis

To build an accurate model of miRNA regulation from the analysis of highly dimensional data such as the one available for the brain of Hdh mice [2], we developed miRAMINT, a pipeline that combines network-based, tree-based and shape-matching analysis into a single workflow (Figure 1) as detailed below.

#### Reduction of data complexity via network analysis

To reduce data complexity, we used WGCNA analysis. To this end, we used the R package WGCNA (https://horvath.genetics.ucla.edu/html/CoexpressionNetwork/Rpackages/WGCNA/). We applied standart settings as previously described [25] to generate signed WGCNA modules from RNA-seq (miRNA and mRNA separately) data in the allelic series of Hdh mice at 2 months, 6 months and 10 months of age, for striatum and cortex, by computing the correlation coefficient across the various CAG repeat lengths. Briefly, we constructed a matrix of pairwise correlations between all gene pairs across condidtions and samples. We removed all genes having less than two counts in all samples. We then constructed a “signed” pairwise gene co-expression similarity matrix and we raised the co-expression similarities to the power β=6 to generate the network adjacency matrix. This procedure removes low correlations that may be due to noise. We then computed consensus modules using maxBlockSize = 500, minModuleSize = 20 and mergeCutHeight = 0.15. The profile of the genes (respectively miRNA) in a module is summarized by the eigen-gene (respectively eigen-mir). To exclude the miRNA modules and mRNA modules that are not correlated, we then computed the Spearman’s score between each possible eigen-mir:eigen-gene pairs. Negative correlations with a false discovery rate lower than 1% using the Benjamini-Hochberg method (Benjamini Y, 1995) were considered statistically significant. This analysis allowed molecular entities that are not correlated at all to be filtered out, based on the lack of negative correlations between eigen-miRNAs and egen-genes.

#### Feature selection

To select the miRNAs that best explain the expression of target genes in the miRNA and mRNA space defined by the paired miRNA:mRNA WGCNA modules, we used RF analysis. Random forests are collections of decision trees that are grown from a subset of the original data. This non parametric method has the advantage of dealing with non-linear effects and of being well-suited to the analysis of data in which the number of variable p is higher than the number of observation. Firstly, we removed the mRNA WGCNA nodes that show no significant deregulation across CAG-repeat lengths and age points. For each target, we then considered all miRNAs in the paired module(s) as possible explanatory variables of the target expression profile across conditions. Then, RF analysis implemented in the R package Ranger was performed by using the Altmann’s approach [26]. This approach has been initially proposed as heuristics in order to correct for the possible bias associated with the traditional measure of variable importance such as the Gini importance measure [26]. This approach has the advantage of using permutation to provides a p-value for the association of each miRNA with a potential target gene, reducing the risk that explanatory variables may be selected by chance. The first step of the Altmann’s approach is to generate an importance score for all variables. Then, the variable to be explained (mRNA) is randomly permutated. Permutation data are then used to grow new random forests and compute the scores for the predictor variables. Permutation were repeated 100 times (default parameter), thus generating 100 scores of importance for each miRNA variable that can be regarded as realizations from the unknown null distribution. These 100 scores were used to compute a p-value for each predictor variable. If the classification error rate for a mRNA was higher than 10%, we rejected the possibility that this mRNA could be under miRNA regulation. When the error rate of classification was lower that 10%, we retained the miRNA(s) associated with mRNA(s) with a p-value < 0.1. Finally, to further ensure the reliability of feature selection, the entire RF analysis, each round recruiting different starting seeds, was repeated until the pool of hypotheses at the intersection of all ensembles of hypotheses generated by all RF iterations is stable. A pool of hypotheses was considered to be stable and RF iterations were stopped when greater than 80% of the hypotheses were conserved across 3 consecutive rounds of analysis. A stable pool of hypotheses was obtained for a range of 3-13 iterations (as illustrated in Figure 1).

#### Shape-matching

The LFCs of a miRNA and a mRNA across multiple conditions (herein as defined by 5 expanded CAG repeat alleles and 3 age points) defines a surface that provides a strong basis for associating a miRNA with its putative target(s). To refine feature selection (see above), we computed the slope of each edge between two conditions. We then computed the Spearman’s score between the slopes for each gene and those for explanatory miRNA(s). Finally, we retained the miRNA-target pairs for which the Spearman’s score is negative and such that the false discovery rate is lower than 0.05 using the Benjamini-Hochberg method (Benjamini Y, 1995).

### Comparison to proteomic data

Previous studies have shown that RNA-seq may validate proteomic data whereas few proteomic data may validate gene deregulation [2]. Nonetheless, we tested whether the deregulation of gene targets retained by MiRAMINT might be also observed at the protein level. To this end, we used the protein data as processed in the HdinHD database (https://www.hdinhd.org/). These data cover 6 CAG-repeat lengths across 3 age points, similarly to RNA-seq data. Briefly, the label-free quantification (LFQ) of the proteins was obtained as previously described [2]. We used the log10 ratio provided in the HDinHD database. This ratio compares the LFQ of the protein for a given CAG repeat length *versus* the LFQ at Q20 for each age. To test for correlation between the deregulation of the mRNA and the deregulation of the protein product, we computed the Spearman’s score between the log-fold-change of the gene and the log10 ratio of the protein. For genes encoding more than one protein in the data-set, we tested for correlation with all protein products and we selected for the one showing the best Spearman’s score. Given the differences in the depthness and dynamics of these data compared to RNA-seq data, a p-value < 0.05 on the Spearman’s score was considered significant.

### Data and code availability

The full list of WGCNA edges that define miRNA and mRNA expression either in the cortex or striatum and a 3D-visualization database of all miRNA-target pairs retained by miRAMINT analysis are available at http://www.broca.inserm.fr/MiRAMINT/index.php. The source code developed for running miRAMINT, written using R, is available http://www.broca.inserm.fr/MiRAMINT/index.php.

## Authors contributions

Design of the study: LM, CN. Data acquisition, analysis and interpretation: LM, SSN, JD, JA, JR, CN. Drafting the manuscript: LM, CN. Conception of the study: CN. All authors read and approved the final manuscript.

## Acknowledgments

We thank Benoit Perthame (Laboratoire Jacques-Louis Lions, Sorbonne Université) for discussions on mathematical modelling. This work was supported by Sorbonne Université, INSERM and CNRS, Paris, France and by the CHDI Foundation (grant number A-12273), Princeton, USA. The funding agencies had no role in study design, data analysis, decision to publish, or preparation of the manuscript.

## Supplemental data

**Table S1.** Lists of nodes in miRNA and mRNA WGCNA modules. Module membership is indicated for mRNAs and miRNAs. NA, not applicable.

**Table S2.** miRNA-target pairs retained by RF analysis. The p-values are the ones provided by the Altmann’s RF algorithm [26]. This table shows the Spearman’s score for surface matching (see Methods) and LFC values on the targets for all miRNA-target pairs.

**Table S3. Surface-matched miRNA-target pairs for which there is evidence for a binding site.** This table shows the miR-target pairs for which (i) the Spearman’s score for plane matching is positive and has a p-value < 0.01 upon Bonferroni correction for multiple testing and (ii) there is evidence for a binding site as supported at least by one database including MicroCosm, TargetScan and miRDB. The LFC amplitude of the target is indicated (see Figure 2 for pairs for which LFC amplitude of the target is above 0.8). The maximum LFC in absolute value of the targets. NA, not applicable.

